# Personality and MB-COMT gene: molecular-genetic and epigenetic associations with NEO-PI-R personality domains and facets in monozygotic twins

**DOI:** 10.1101/2024.05.20.594935

**Authors:** Dušanka Mitrović, Snežana Smederevac, Lissette Delgado-Cruzata, Selka Sadiković, Dejan Pajić, Mechthild Prinz, Zoran Budimlija, Milan Oljača, Jelena Kušić-Tišma, Nataša Vučinić, Aleksandra Milutinović

**Affiliations:** Department of Psychology, Faculty of Philosophy, University of Novi Sad, Serbia Dr Zoran Djindjic 2 21000 Novi Sad, Serbia; John Jay College of Criminal Justice, City University of New York, New York, NY 524 West 59th. Street, New York, NY, 10019, United States; Department of Neurology, NYU School of Medicine, New York, NY 240 East 38th Street, New York, NY 10016, United States; Institute of Molecular Genetics and Genetic Engineering (IMGGE), University of Belgrade Vojvode Stepe 444a, 11042 Belgrade 152, Serbia; Faculty of Medicine, University of Novi Sad, Serbia; Hajduk Veljkova 3 21000 Novi Sad, Serbia

**Keywords:** MB-COMT, DNA methylation (DNAm), NEO-PI-R, monozygotic twins

## Abstract

This study investigates the relationship between *MB-COMT* DNA methylation (DNAm) and the personality traits outlined in the NEO-PI-R model through an epigenetic study of monozygotic twins. DNAm, a critical epigenetic mechanism, regulates gene expression and has been linked to various biological processes and disorders. By leveraging the genetic similarities of monozygotic twins, this research explores how epigenetic variations influenced by environmental factors correlate with personality differences. The study utilized the Five-Factor Model (FFM) to categorize personality traits into five domains: Neuroticism, Extraversion, Conscientiousness, Agreeableness, and Openness to Experience. Each domain comprises six facets, providing a granular view of personality. The research centered on the catechol-O-methyltransferase (*COMT*) gene, focusing on its role in dopamine metabolism, which is hypothesized to influence personality traits through the dopaminergic system. DNAm status in the promoter region of the *MB-COMT* gene, was examined to determine its association with personality facets. Preliminary findings suggest a complex interaction between *MB-COMT* DNAm patterns and personality traits. Specific methylation patterns at different CpG sites were linked to varying expressions of traits such as impulsivity and aggression, highlighting the nuanced impact of epigenetics on personality. This study underscores the potential of integrating genetic, epigenetic, and environmental data to enhance our understanding of personality formation. The results contribute to a broader understanding of how genetic predispositions shaped by environmental factors manifest in complex trait differences, paving the way for future research in genetic psychiatry and personalized medicine.

## Introduction

Epigenetic modifications, notably DNA methylation (DNAm), are pivotal in regulating gene expression. The prominence of DNAm within regulatory domains such as gene promoters is widely recognized, primarily due to its association with the suppression of gene expression, as elucidated by Jones (1). This epigenetic mechanism is instrumental in orchestrating a myriad of biological processes and plays a significant role in the emergence and manifestation of various disorders. Contemporary research has primarily focused on exploring the impact of DNA methylation in the context of personality and mental disorders. This research trajectory is driven by the hypothesis that these disorders often display more conspicuous and distinguishable phenotypic characteristics, as noted by Gescher et al. (2) and Thomas et al. (3). Comprehending the epigenetic underpinnings of such disorders is imperative for deciphering their complex etiology and potentially developing targeted therapeutic strategies.

However, investigating the influence of environmental factors on epigenetic variations presents a distinct challenge, particularly when examining personality traits. In contrast to disorders, personality traits signify dimensions of individual differences that typically demonstrate consistency in their expression and intensity over time. To elucidate the epigenetic foundations of these traits, a multifaceted approach is required, encompassing various levels of the trait hierarchy.

In this context, twin studies represent an invaluable resource. These studies leverage the genetic congruence of monozygotic twins (MZ), who share nearly 100% of their genetic makeup, to scrutinize disparities in their epigenome and phenotype. By contrasting MZ twins with divergent epigenetic profiles, researchers can discern the influence of environmental factors on epigenetic alterations and their correlations with personality traits.

### The Five-Factor Model: A Genetic and Environmental Perspective

The Five-Factor Model (FFM) is widely acknowledged in the psychological literature as a relevant framework for personality analysis. The model comprises five dimensions - Neuroticism, Extraversion, Conscientiousness, Agreeableness, and Openness to Experience, called personality domains, with each domain encompassing six lower-order facets. The best-known operationalization of this model is the NEO-PI-R questionnaire, an extensively validated and reliable tool for personality assessment (4–6), widely recognized for its universality. The stability of the FFM has been attributed in part to the genetic basis underlying traits and shaping their domains (7), a proposition further supported by studies highlighting significant genetic influences on personality traits (8,9). However, some research (10–12) suggests that the genetic and environmental architecture of personality traits may deviate from their phenotypic manifestations, with individual variances being more influenced by specific genetic and environmental factors than common ones.

The hierarchical structure of the FFM, especially its lower levels of hierarchy, such as facets and nuances, has attracted research attention (13). These facets, representing more subtle and specific phenotypic expressions within each domain, have shown incremental validity and are influenced by unique genetic and environmental factors (14–16). Notably, the reliance only on aggregate scores for each dimension of the FFM is acknowledged as a potential limitation, as it may reduce the statistical power of research in this area (10).

Twin studies play a key role in examining the genetic contribution to individual differences in personality traits, indicating that heritability accounts for approximately 50% of the variance (11,17–20). However, variability in heritability estimates across studies suggests the presence of non-additive gene effects and gene interactions. High genetic correlations among personality traits are hypothesized to result from pleiotropy (12), linking trait similarity between individuals to their genetic resemblance (e.g. 21).

Examining the complex interaction between genetic effects and environmental factors, it becomes apparent that the same gene variant can produce different phenotypic outcomes, and different combinations of genes can result in identical phenotypes. This complexity underlines the importance of identifying small, consistent gene variant effects on personality traits (22). Since genes indirectly influence psychological phenomena through biochemical and physiological processes, consideration of brain structure and function appears as a suitable endophenotype for tracing the path from genes to behavior (23,24).

### The Dopaminergic System and Its Influence on the FFM

The dopaminergic system, predominantly expressed in the various brain regions, plays a vital role in dopamine biosynthesis and synaptic regulation (25–29). Existing research, despite presenting mixed findings, generally associates dopaminergic genes with various personality traits, including approach behaviors, extraversion, particularly positive emotionality (29–31), impulsivity (32), openness to experience (26), and neuroticism, primarily through anxiety, depression, and harm avoidance (25,33,34), as well as the behavioral activation system (17). The dopaminergic hypothesis of extraversion suggests a link between the COMT enzyme, critical for dopamine metabolism, and extraversion, either independently (35) or in interaction with other dopamine-related genes (36).

The catechol-O-methyltransferase (*COMT*) gene, located on chromosome 22q11, is responsible for encoding the COMT enzyme, a pivotal agent in the primary inactivation pathway of dopamine within the brain, particularly manifesting substantial expression in the prefrontal cortex (25). This enzyme plays a crucial role in the metabolic decomposition of catecholamines, including adrenaline, noradrenaline, and dopamine (25,30,31), thereby exerting a significant influence on a spectrum of cognitive functions and personality attributes (25,27). A notable functional polymorphism of the *COMT* gene involves the substitution of valine (Val) for methionine (Met) at codon 158, which consequently alters the enzyme’s thermal stability and its enzymatic activity (27,31,37). This modification results in a reduction of enzyme activity by approximately 30-35% in the prefrontal cortex of individuals carrying the Met allele (25), leading to an amplification of dopamine signaling due to the predominant role of COMT in dopamine clearance from synaptic spaces (27,30).

Previous research has demonstrated a correlation between neuroticism and diminished activity of the COMT enzyme (25), with a notable gender-specific manifestation. The connection with anxiety-related traits, for instance, was predominantly observed in women (34). Conversely, in men, the Val allele has been linked to increased negative emotionality (25), neuroticism, and harm avoidance tendencies (38). Furthermore, significant phenotypic variances between carriers of the Val and Met alleles of the *COMT* gene have been documented in relation to extraversion (31), positive emotionality (25), and novelty-seeking behaviors (30). Notably, Met/Met homozygotes have demonstrated augmented prefrontal cortical activity during phases of reward anticipation (39), suggesting that the directional association, akin to that with neuroticism, may exhibit gender-specific divergences: women with the Met allele showing reduced extraversion (31), and men with the Val allele demonstrating the opposite (30).

Moreover, the *COMT* gene has been implicated in the realm of creative potential (40), where a decrease in dopamine removal correlates with elevated synaptic dopamine levels, potentially contributing to enhanced openness traits (27,28).

### *MB-COMT* DNA Methylation and Endophenotypes

The catechol-O-methyltransferase (*COMT*) gene occupies a central role in the metabolic processing of catecholamines within the brain, including neurotransmitters such as dopamine. Of particular significance is the membrane-bound catechol-O-methyltransferase (*MB-COMT*), a notable isoform of this gene. Elucidation of the epigenetic regulation of the *MB-COMT* gene promises to shed light on the molecular determinants underpinning individual variances in behavioral, cognitive, and emotional regulatory mechanisms. Attributes such as impulsivity, anxiety, and cognitive capabilities, notably working memory and executive control, are postulated as potential endophenotypes susceptible to modulation by *MB-COMT* DNAm.

Recent scholarly endeavors have delved into the role of DNAm of the *MB-COMT* gene in the modulation of personality traits, with a particular focus on impulsivity. It has been observed that individuals exhibiting specific DNAm patterns in the *COMT* gene may display varied capacities in modulating impulsive behavior. Dopamine, crucially modulated by COMT, is integral to decision-making and impulse control processes. Variations in *COMT* DNAm are hypothesized to influence the efficacy of dopamine degradation, thereby contributing to interindividual differences in impulsivity (41). Furthermore, investigations into the correlation between *MB-COMT* DNAm and personality traits, guided by the Revised Reinforcement Sensitivity Theory, have unveiled insights into how *MB-COMT* gene DNAm may affect behavioral patterns and personality traits. This is particularly evident in traits associated with responses to environmental stimuli, such as impulsivity and aggression (41).

### Current Study

This study seeks to contribute to a comprehensive understanding of the interactions between genes, environment and personality traits by integrating genetic, epigenetic and phenotypic data. By such an approach, we aim to advance the insight into the underlying mechanisms shaping personality.

Specifically, our research will explore the relationship between variations in the catechol-O-methyltransferase (*COMT*) gene and personality traits within a five-factor model. Furthermore, we will investigate how differences in *MB-COMT* promoter DNAm levels correlate with personality traits. Our approach involves utilizing twin data, relying on the unique genetic similarity inherent in monozygotic twins that allows for discerning the epigenetic changes resulting from environmental influences. Thus, our study aims to assess both genetic and environmental contributions to individual variation in personality traits, with a particular focus on the hierarchical structure of the FFM, including its facets.

## Method

### Procedures and Participants

The comprehensive recruitment, testing, and data collection process within the Serbian Advanced Twin Registry (STAR) is detailed in Smederevac et al. (42). From the STAR, which includes data on 1654 participants (827 twin pairs), we selected all monozygotic twin pairs that had data on all relevant phenotypic measures and were genotyped for *MB-COMT* (*rs4680*). The study obtained the required ethical clearances from the Institutional Ethical Committees, with the codes #02-374/15, #01-39/229/1, and #O-EO-024/2020. Participation in the study was entirely voluntary, with all participants providing informed written consent before their involvement.

#### Single Nucleotide Polymorphism (SNP) Sample

After excluding certain cases due to failed genotyping, the sample comprised 430 twins. The age of the participants ranged from 18 to 60 years (M = 24.66; SD = 7.72; 74.4% women). The sample included more monozygotic (75.8%) than dizygotic twins. Monozygotic twins were aged between 18 to 60 years (M = 25.18; SD = 8.08; 75.5% women). Dizygotic twins ranged from 18 to 48 years of age (M = 23.05; SD = 6.21; 71.2% women).

#### Epigenetic Sample

To investigate the association between DNAm differences and behavioral phenotypic traits, we analyzed a subset of 70 monozygotic twins (MZ) from the SNP sample, who had high-quality buccal swab DNA suitable for methylation studies. This subset included 16 male and 54 female twins, aged between 18 and 44 years, with an average age of 23.30 years (SD = 6.68).

### Phenotypic Measures

Revised NEO-PI-R Personality Inventory (4,43). The NEO-PI-R personality inventory contains 240 items with a five-point Likert response format and is designed to operationalize the five major personality dimensions of the five-factor model: Neuroticism (N), Extraversion (E), Agreeableness (A), Conscientiousness (C), and Openness to experience (O). Each dimension includes 6 lower-order facets (30 in total), operationalized through 8 items for each personality aspect. Reliabilities (α) range from .30 for Openness to .78 for Neuroticism.

### Zygosity Analysis

The zygosity of the subjects was determined by DNA isolated from a buccal swab sample by analyzing microsatellite loci. The Investigator24plex GO! kits were used for microsatellite analysis. (Qiagen®, Valencia, CA, USA) or (Applied Biosystems®, Thermofisher Scientific, Waltham, MA, USA) according to the manufacturer’s instructions. Buccal swabs are tested using STR (short tandem repeat) megaplex sets according to the manufacturer’s instructions, and they are of two types: Investigator 24plex GO! (Qiagen®, Valencia, CA, USA) and GlobalFiler (Applied Biosystems®, Thermofisher Scientific, Waltham, MA, USA).

Both kits detect 21 autosomal microsatellite loci. Samples with partial profiles were interpreted if a result was present at at least 10 gene loci. Fully concordant (identical) pairs are categorized as monozygotic twins based on microsatellite profiles, and all others as dizygotic twins. One part of the DNA samples was analyzed at the Institute of Forensic Medicine in Novi Sad, and the other at the John Jay College of Criminal Law in New York.

#### The Genotyping of COMT (rs4680) Polymorphisms

The *COMT* gene (rs4680) genotyping was performed using the TaqMan assay (TaqMan SNP, Applied Biosystems®, Warrington, UK) at the Faculty of Medicine, University of Novi Sad. TaqMan single nucleotide polymorphism genotyping assays are based on 5’-nuclease activity to detect and amplify specific polymorphisms in purified DNA samples and use probes that bind to the minor groove for better discrimination and accurate determination of allelic type. Samples for polymerase chain reaction (PCR) were prepared from 10 ng of genomic DNA with 1 µl of TaqMan genotyping assay and 12.5 µl of genotyping master mix in a final volume of 25 µl. For the polymerase chain reaction, a 96-well microplate, and an ABI Prism 7500 Fast PCR device (Applied Biosystems®, Foster City, California, USA) are used.

The *COMT rs4680* gene was defined by three genotypes: 129 highly active homozygotes (Met/Met), then 224 heterozygotes (Met/Val), and 77 low active homozygotes (Val/Val). Individuals with at least one copy of the Val allele were grouped into the Val group (301 subjects), while Met homozygotes formed the Met+ group (129 subjects). *COMT rs4680* gene was in Hardy-Weinberg equilibrium (χ2 = 0.85; *df* = 2; *p* > .05).

The *COMT* genotypes in the epigenetic sample were also categorized into the Met/Met group (23 subjects), Met/Val group (37 subjects), or Val/Val group (10 subjects). All individuals who had at least one copy of the Val allele were grouped into the Val group (47 subjects), while Met homozygotes formed the Met+ group (23 subjects). The *COMT rs4680* gene in the epigenetic sample was in HWE (χ2 = 5.06; *df* = 2; *p* > .05).

### DNA Methylation Analysis

Genomic DNA was isolated from buccal swabs utilizing the QIAamp DNA Mini Kit (Qiagen®, Valencia, CA, USA), adhering to the protocol provided by the manufacturer. The DNA underwent bisulfite conversion through the EZ DNA Methylation-Gold kit (Zymo Research), following its specific instructions. The bisulfite-treated DNA was then eluted in 15μl of water, and an aliquot of 2.5-5μl was employed for Pyromark PCR amplification (Qiagen). This assay was designed to target a 228-bp segment within the MB-COMT promoter region. The forward primer sequence was 5’-TGGGGTAGATTAGGGTTGT-3’, and the biotinylated reverse primer at the 5 ’end was 5’-CCACACCCTATCCCAATATTC-3’. The amplification process followed the guidelines of the Pyromark PCR Kit (Qiagen). Pyrosequencing was conducted on a PyroMark Q24 system (Qiagen), adhering to the manufacturer’s protocol. The sequencing primer used was 5’-GGATAGGGGAGGGTTTAGTT-3’, and the sequence analyzed was 5’-TYGGGYGGGTYGTYGYGGGAGAGGTGAGAG-3’. This method assessed the DNAm levels of five CpG sites. DNAm was analyzed using PyroMark Q24 Advanced 3.0.0 software. The results were expressed as average percent DNAm across the five CpG sites, or specific DNAm levels at individual CpG sites, as indicated in each analysis. Each amplification and pyrosequencing batch included fully methylated and unmethylated DNA (Zymo Research) as controls, alongside no-template controls in all runs.

### Data Analysis

#### Data analysis on the SNP dataset

The statistical analyses applied to the SNP dataset encompassed descriptive statistics, correlations, t-tests, and non-parametric alternatives to the t-test for independent samples, post-hoc tests, and effect size estimation. All analyses were conducted using the JASP statistical program (44). The level of statistical significance for each of the applied t-tests was adjusted using the Bonferroni correction for multiple comparisons. In cases where certain prerequisites for using the t-test were not met (conditions of homogeneity of variances or normality of distribution), alternatives to the t-test for independent samples were employed, namely Welch’s test (for heterogeneous variances) or the non-parametric Mann-Whitney U test (for distributions significantly deviating from normal). For Welch’s test, Cohen’s d was presented as the effect size, and for the Mann-Whitney U test, rank biserial correlation coefficients were used. The common interpretation of effect size for Cohen’s d is small (d = .20), medium (d = .50), or large (d = .80), based on the guidelines provided by Cohen (45). Along with the effect size, confidence intervals for the effect size (95% CI) were provided.

#### Data analysis on the epigenetic dataset

The primary goal of this analysis was to detect association strength between DNAm levels at various CpG sites and the levels of personality traits. Due to high skewness and large variations in value ranges (see the Results section), twin pairs were compared dichotomously. They were divided into four categories based on two criteria: higher/lower level of methylation and higher/lower level of trait expression. For instance, the “lower-lower” group included twin pairs where the first twin exhibited both a lower methylation level at a specific CpG site and a lower score on a personality trait or facet.

Two-by-two contingency tables for each pair of CpG site and trait/facet were analyzed using Barnard’s unconditional exact test. This test was selected over the more commonly used Fisher’s exact test because the goal was to determine whether higher levels of methylation are associated with higher or lower levels of trait expression. Consequently, Barnard’s test was expected to be more appropriate since it does not condition on both margins being fixed, allowing for potentially greater test power in detecting associations. This gain in power is particularly noticeable when sample sizes are small and research scenarios imply multiple comparisons (46).

All statistical analyses in this section were performed using the Python SciPy library (47). Contingency tables were visualized as bubble charts using the Python Plotly library (48).

## Results

### Main effects of the *COMT (rs4680)* gene

In Table A in Supplementary material the descriptive statistics for the FFM domains and facets of neuroticism, extraversion, openness to experience and conscientiousness on groups of *COMT* gene (*rs4680*) carriers were given. The results indicate significant and robust effects of the *COMT rs4680* genotype on self-consciousness from the domain of neuroticism (U = 22341.5; p < .05; p_bonf_ < .05; d = .151; d(CI) = .033-.265). Those in the Met+ group show significantly higher scores on self-consciousness than those in the Val+ group (Table A).

The *COMT rs4680* genotype shows robust main effects on extraversion (U = 15777.5; p < .01; p_bonf_ < .01) and facets of extraversion: warmth (U = 15371.5; p < .01; p_bonf_ < .01), gregariousness (U = 15564; p < .01; p_bonf_ < .01) and positive emotions (U = 16275.5; p < .01; p_bonf_ < .05) (Table 1). The main effects of the *COMT rs4680* genotype on the facets of extraversion remain significant after correction for multiple comparisons, and effect sizes are largest for warmth (mean effect size; d = -.208; d(CI) = -.319- -.092), gregariousness (d = - .198; d(CI) = -.310- -.082), and extraversion domain (d = -.187; d(CI) = -.299- -.070), and Val + group of alleles show statistically significantly higher scores on extraversion, warmth, gregariousness, and positive emotions (Table 1).

**Table 1.** Main effects of the COMT rs4680 genotype on domains and facets of neuroticism, extraversion, openness to experience, and conscientiousness.

|  | *t/U* | *df* | *p* | *p<sub>bonf</sub>* | *d\** | 95% *IP* |  |
| --- | --- | --- | --- | --- | --- | --- | --- |
|  |  |  |  |  |  | *higher* | *lower* |
| Neuroticism (trait) | 21691 |  | .054 | .108 | .117 | -.001 | .233 |
| Anxiety | 21007 |  | .176 | .352 | .082 | -.037 | .199 |
| Hostility | 1.34 | 428 | .181 | .362 | .141 | -.065 | .347 |
| Depression | 21066.5 |  | .159 | .318 | .085 | -.034 | .202 |
| **Self-consciousness** | **22341.5** |  | **.013** | **.026** | **.151** | **.033** | **.265** |
| Impulsiveness | 18854 |  | .634 | 1.00 | -.029 | -.147 | .090 |
| Vulnerability | 21495 |  | .077 | .154 | .107 | -.012 | .223 |
| **Extraversion (trait)** | **15777.5** |  | **.002** | **.004** | **-.187** | **-.299** | **-.070** |
| **Warmth** | **15371.5** |  | **.000** | **.000** | **-.208** | **-.319** | **-.092** |
| **Gregariousness** | **15564** |  | **.001** | **.002** | **-.198** | **-.310** | **-.082** |
| Assertiveness | 19012 |  | .732 | 1.00 | -.021 | -.139 | .098 |
| Activity | 17014 |  | .041 | .082 | -.124 | -.239 | -.005 |
| Excitement seeking | 18979.5 |  | .712 | 1.00 | -.022 | -.141 | .097 |
| **Positive emotions** | **16275.5** |  | **.008** | **.016** | **-.162** | **-.275** | **-.044** |
| Openness to experience (trait) | 20341 |  | .432 | .864 | .048 | -.071 | .165 |
| Fantasy | 20450.5 |  | .377 | .754 | .053 | -.066 | .171 |
| Aesthetics | 20917 |  | .201 | .402 | .077 | -.042 | .194 |
| Feelings | 19397 |  | .988 | 1.00 | -.001 | -.120 | .118 |
| Actions | 17727.5 |  | .150 | .300 | -.087 | -.203 | .032 |
| Ideas | 20170 |  | .520 | 1.00 | .039 | -.080 | .157 |
| Values | 19790 |  | .748 | 1.00 | .019 | -.100 | .138 |
| Conscientiousness (trait) | 17862 |  | .189 | .378 | -0.08 | -.197 | .039 |
| **Competence** | **15971.5** |  | **.003** | **.006** | **-.177** | **-.290** | **-.060** |
| Order | 20584.5 |  | .321 | .642 | .060 | -.059 | .178 |
| Dutifulness | 19303.5 |  | .925 | 1.00 | -.006 | -.124 | .113 |
| **Achievement striving** | **16599** |  | **.017** | **.034** | **-.145** | **-.259** | **-.027** |
| Self-discipline | 18194.5 |  | .299 | .598 | -.063 | -.180 | .056 |
| Deliberation | 18021.5 |  | .236 | .472 | -.072 | -.189 | .047 |

*Notes.* *t/U* - in relation to the fulfilled conditions for the application of the t-test (normality of distribution and homogeneity of variances), the column *t/U* contains the result of the t-test for independent samples, the Welsh test (in case of inhomogeneity of variances) and Mann-Whitney U-test in case of violation of normality of distributions; *df* – degrees of freedom are given for the t-test; \* - results calculated using the Welsh test are given under an asterisk; *p<sub>bonf</sub>* – significance level corrected for multiple comparisons; *d\** - effect size given as Cohen's *d* for t-test and Welsh test; in the case of the Mann-Whitney test, the rank of the biserial correlation was calculated as the size of the effect; 95% *IP* - confidence interval of the effect size.

Moreover, the *COMT rs4680* genotype shows significant main effects on competence (U = 15971.5; p < .01; p_bonf_ < .01; d = -.177; d(CI) = -.290- -.060) and achievement striving (U = 16599; p < .05; p_bonf_ < .05; d = -.145; d(CI) = -.259- -.027). As with the extraversion domain, the carriers of the Val+ allele show statistically significantly higher scores on both the competence and the achievement striving. The results indicated that the *COMT* rs*4680* genotype showed no significant main effects on other aspects of neuroticism, extraversion, openness to experience and conscientiousness.

In Table B of the Supplementary Material, the main effects of *COMT rs4680* genotype status on *MB-COMT* promoter CpG levels are presented. There were no statistically significant effects of the *rs4680* genotype on the DNAm levels of any CpGs.

### Association between DNAm differences and behavioral phenotypic traits

Table 2 presents basic descriptive statistics for DNAm levels at five analyzed CpG sites and scores across five NEO-PI-R domains. Due to a violated assumption of normality for DNAm levels, we opted for Barnard’s test as a more robust non-parametric method for assessing the association between categorical variables. Another rationale for this choice was the intention to assess association strength in terms of contingencies, specifically to compare twins within each pair and evaluate the likelihood of a twin having higher level of methylation to also obtain higher or lower score on a specific personality domain or facet. In line with that, we divided twin pairs into four groups based on two dichotomous criteria: higher/lower score on DNAm level for the first twin and higher/lower score on NEO-PI-R domain or facet for the same twin in a twin pair.

**Table 2.** Descriptive statistics for DNAm levels on five CpG sites and scores on five main NEO-PI-R domains.

|  | M | Mdn | s | Min | Max | Sk | Ku |
| --- | --- | --- | --- | --- | --- | --- | --- |
| CpG1 | 7.75 | 5.77 | 6.43 | 0 | 33.74 | 1.82 | 4.04 |
| CpG2 | 4.55 | 3.43 | 4.58 | 0 | 23.85 | 2.39 | 6.30 |
| CpG3 | 6.18 | 4.34 | 5.47 | 0 | 25.90 | 1.73 | 2.96 |
| CpG4 | 6.16 | 4.28 | 5.21 | 0 | 26.50 | 1.87 | 3.53 |
| CpG5 | 4.40 | 3.11 | 4.22 | 0 | 22.41 | 2.01 | 4.50 |
| Neuroticism | 87.31 | 90.00 | 20.21 | 40 | 120 | -0.33 | -0.71 |
| Extraversion | 118.04 | 117.50 | 17.67 | 72 | 159 | 0.10 | -0.28 |
| Openness | 114.60 | 112.00 | 14.96 | 83 | 156 | 0.31 | -0.07 |
| Agreeableness | 115.04 | 116.50 | 14.34 | 67 | 141 | -0.73 | 1.04 |
| Conscientiousness | 122.26 | 122.50 | 20.13 | 68 | 157 | -0.44 | -0.24 |

Notes: M – Mean, Mdn – Median, s – Standard deviation, Min/Max – Minimal and maximal values, Sk – Skewness, Ku - Kurtosis

Hypothesis testing was conducted using a one-sided approach. Let p_1_ represent the theoretical binomial probability that a twin has a higher DNAm level and a lower score on a personality trait compared to his or her sibling. Let p_2_ denote the theoretical binomial probability that a twin has both a higher DNAm level and a higher score on a personality trait than his or her sibling. Setting the alternative (tail) to “greater” would then test the assumption: H_0_: p_1_ ≤ p_2_ vs. H_1_: p_1_ > p_2_. Conversely, setting the alternative (tail) to “less” would test the assumption: H_0_: p_1_ ≥ p_2_ vs. H_1_: p_1_ < p_2_. In line with these hypotheses, lower score categories were treated as “not higher” under the “greater” alternative, and hence included pairs of twins with identical scores on the personality trait/facet. Similarly, higher score categories were considered “not lower” under the “less” alternative, thus including pairs of twins who had the same score on the personality trait/facet. Only one twin pair had identical DNAm levels (zero) at only one site (CpG2), and this pair was excluded from all the analyses that included this CpG site. Tables C and D of the Supplementary Materials show Barnard’s test values for association between each DNAm level and each NEO-PI-R dimension and facet.

Figure 1 shows contingency tables, visualized as bubble charts, and corresponding Barnard’s test values for statistically significant associations between methylation levels on five CpG sites and NEO-PI-R facets under the “greater” alternative hypothesis. In general, higher DNAm levels are associated with higher scores on facets of Extraversion (Gregariousness and Warmth) and Agreeableness (Trust and Modesty). Additionally, Values facet has a bordering significant association with the DNAm level on CpG3 site.

**Figure 1.**
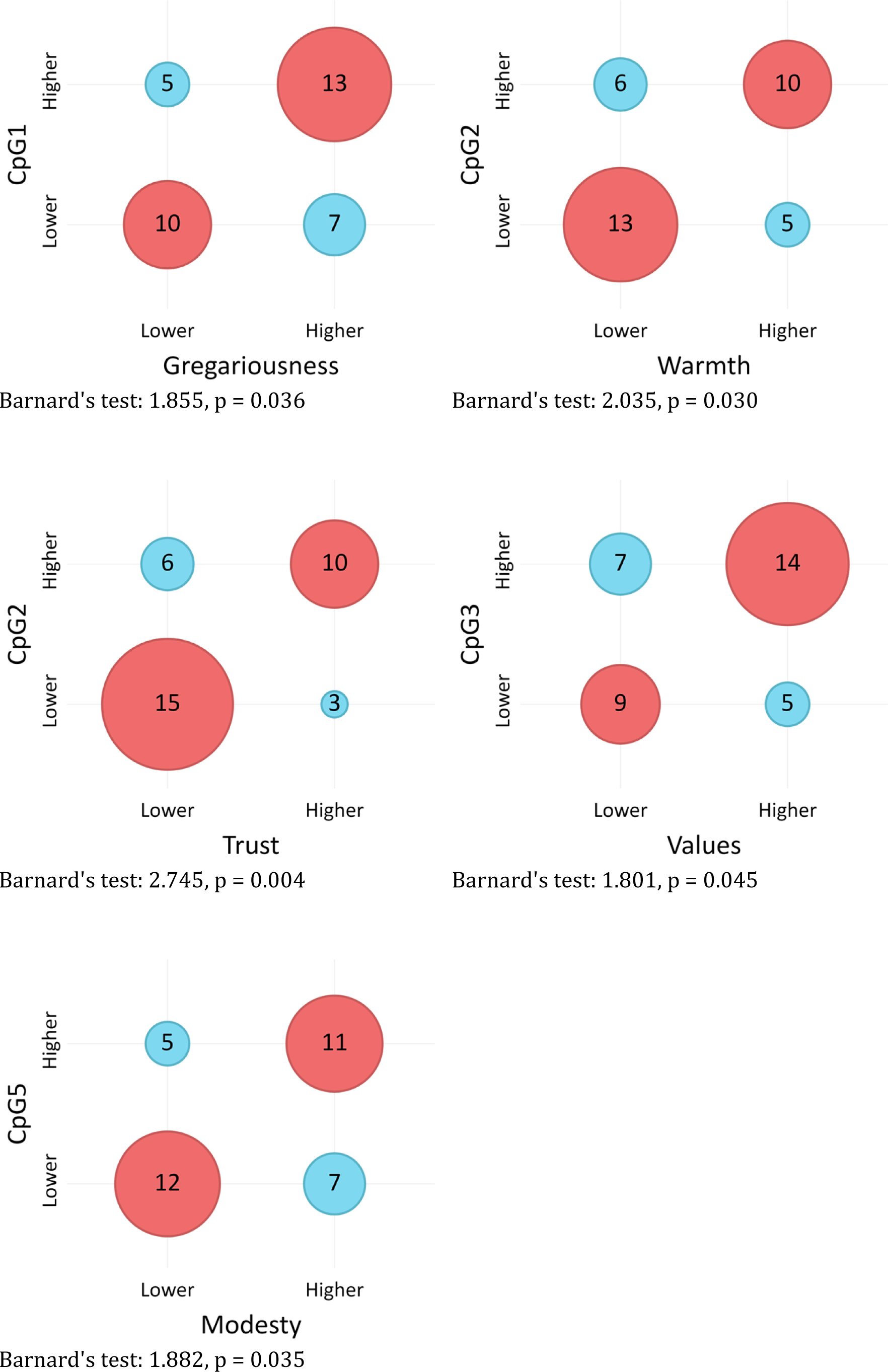
*Association between COMT DNAm levels and NEO-PI-R facets (“greater” alternative)*

Figure 2 shows contingency tables and corresponding Barnard’s test values for statistically significant associations between DNAm levels on five CpG sites and NEO-PI-R traits and facets under the “less” alternative hypothesis. Lower DNAm levels on multiple CpG sites are primarily associated with higher scores on facets of Neuroticism (Hostility, Depression, Impulsiveness). Extraversion and its facets (Excitement Seeking, Positive Emotion), as well as Agreeableness and its facets (Straightforwardness, Tender-Mindedness) again show significant associations, but this time in opposite directions and at different CpG sites.

**Figure 2.**
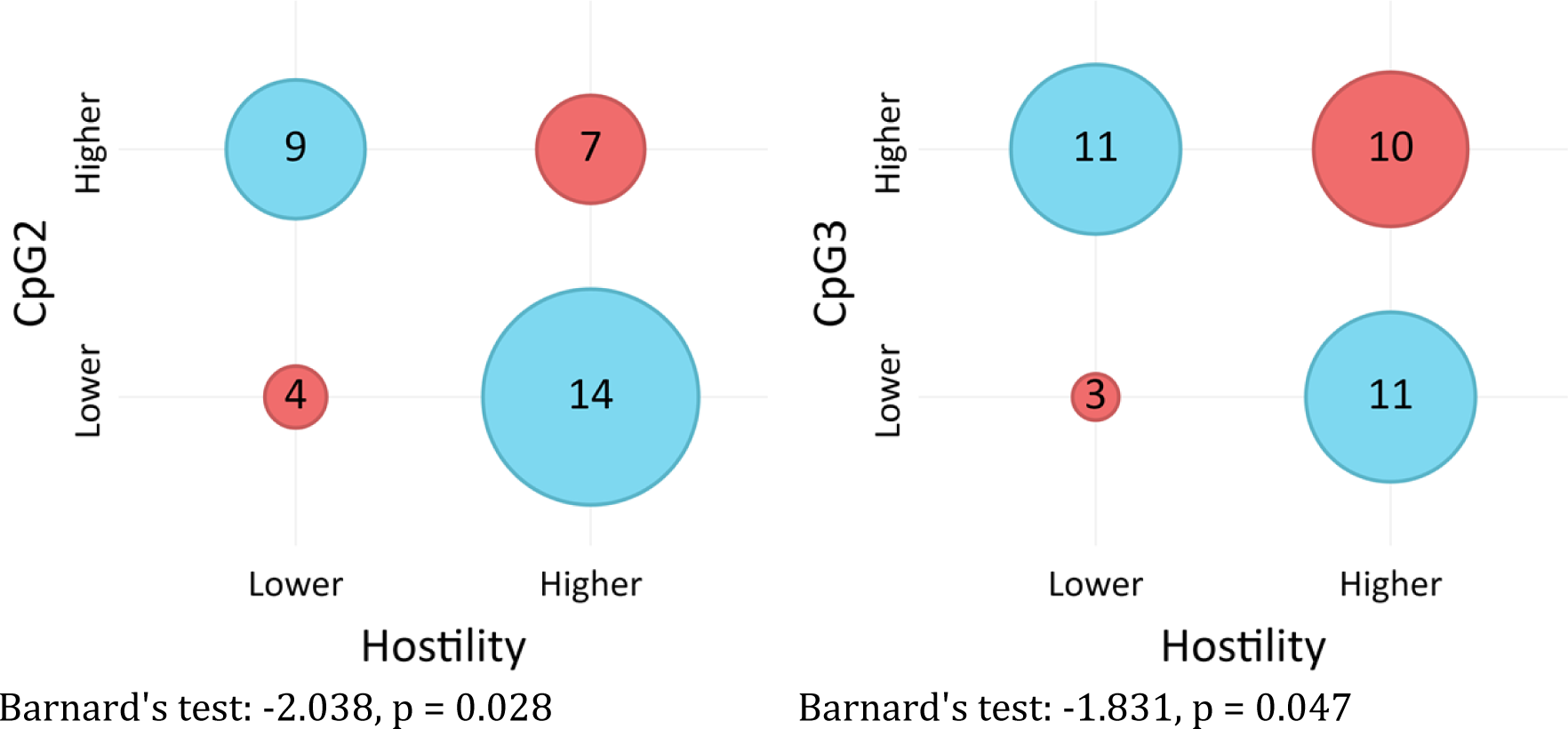
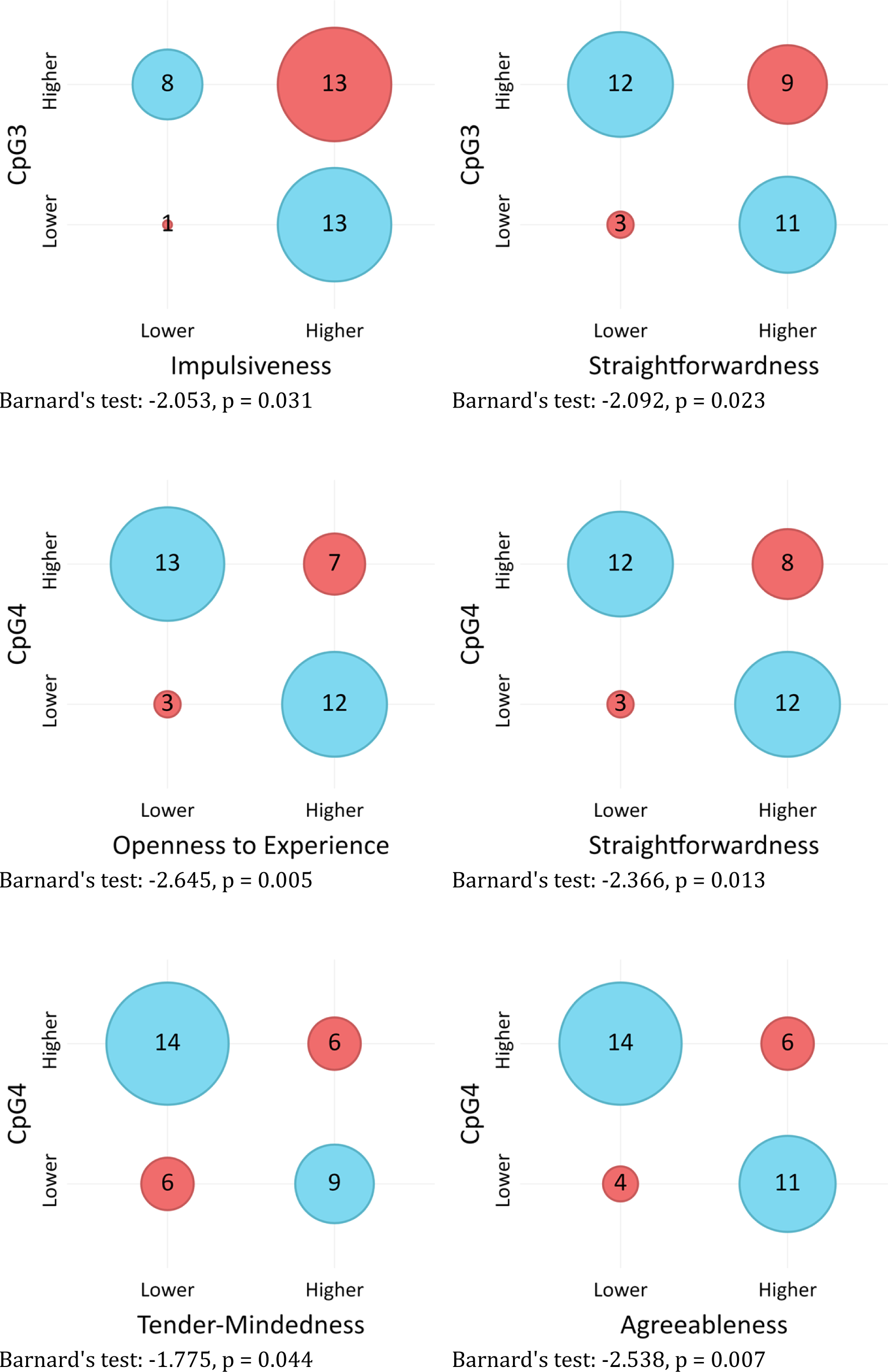
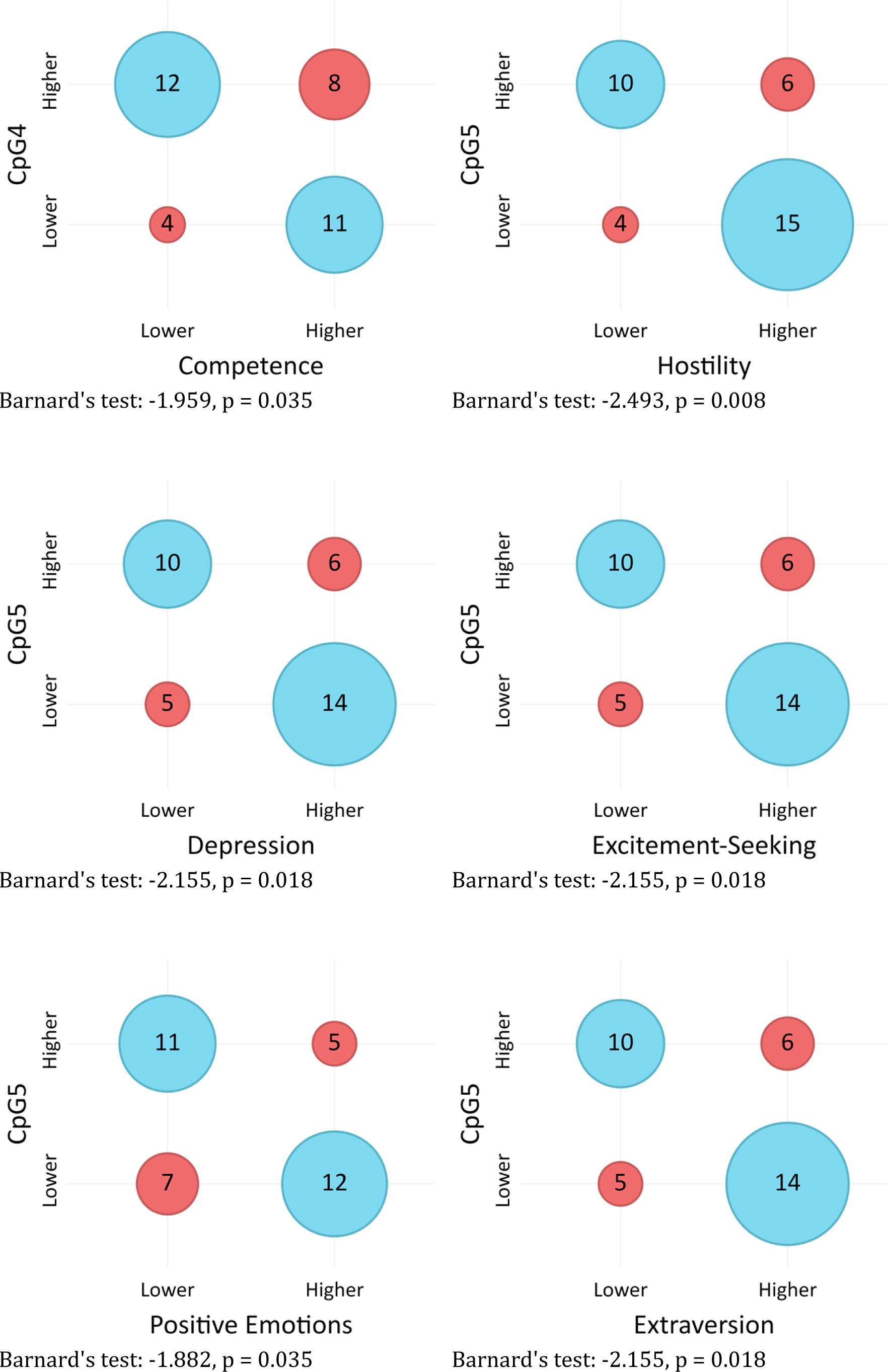
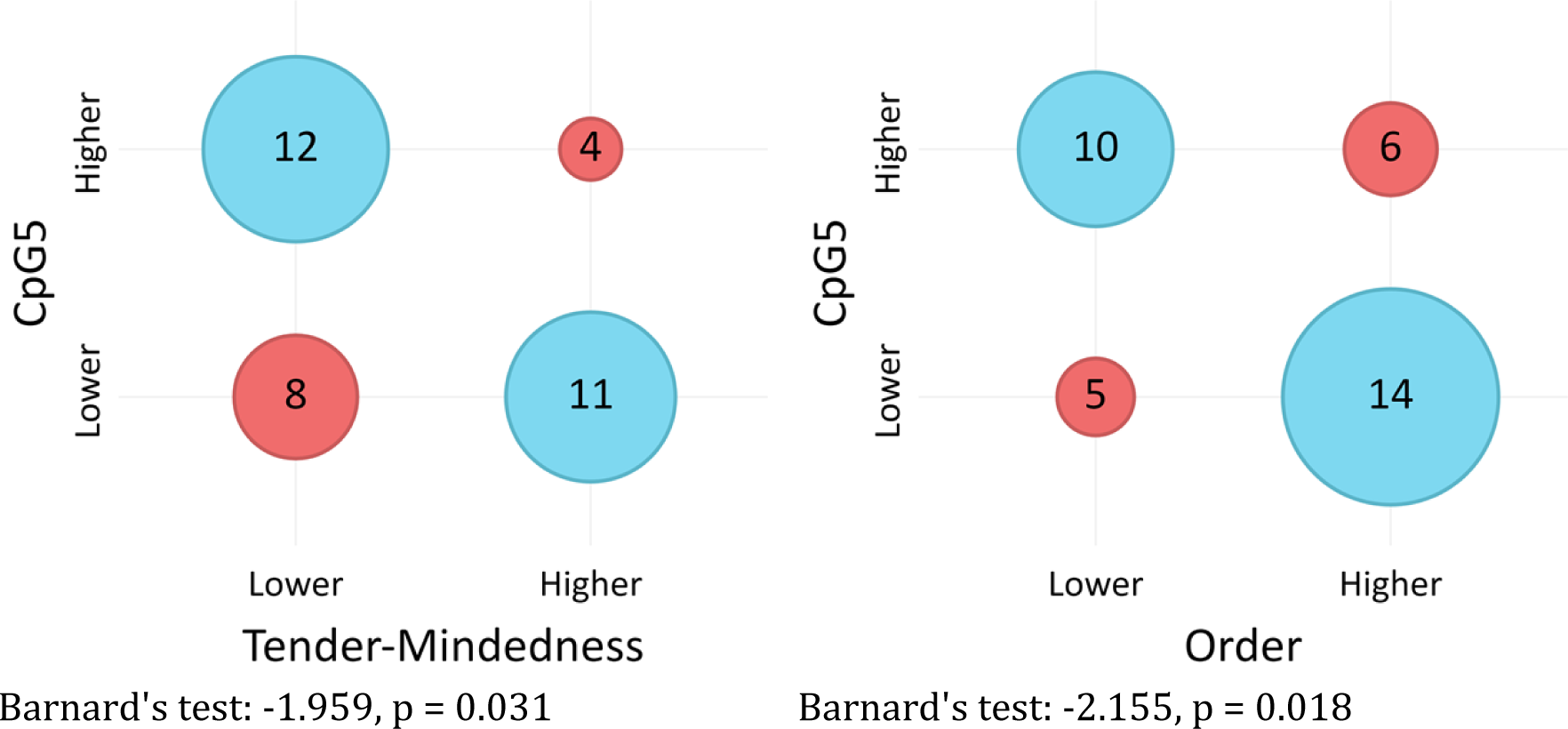
*Association between COMT DNAm levels and NEO-PI-R facets (“less” alternative)*

## Discussion

The main goal of this research was to examine the role of the *COMT rs4680* genotype in the genetic basis of personality traits according to the five-factor model. In addition to the relationships between the polymorphism of this gene and the domains and facets of the FFM, we were also interested in whether the level of DNAm of the promoter region of this gene, assumed to be influenced by environmental factors, shows a connection with different levels of FFM domains and facets.

### The Met158Val polymorphism of the *COMT* gene and personality traits

Results of the association study indicate a connection between the *COMT* gene *rs4680* polymorphism and traits within the domains of Neuroticism, Extraversion, and Conscientiousness. Specifically, individuals carrying the Met allele tend to exhibit heightened levels of Self-Consciousness (N), whereas those with the Val allele display higher Extraversion, as well as its facets Gregariousness, Warmth, and Positive Emotions. They also score higher on competence and Achievement Striving within the Conscientiousness domain.

Thus, reduced *COMT* enzyme activity in Met allele carriers, resulting in higher synaptic dopamine levels (30), is linked to social anxiety and heightened sensitivity to social reinforcement. Due to increased dopaminergic signaling, individuals carrying the Met allele may experience greater self-consciousness and display more inhibited behavior in social interactions, as dopamine is associated with behavioral control (49). Heightened reactivity and focus on internal experiences could potentially result in a tendency to interpret social cues as threatening.

Conversely, heightened *COMT* enzyme activity in Val allele carriers, resulting in reduced dopaminergic activity, is associated with sociability, friendly behavior, and a positive mood, aligning with the dopamine hypothesis of extraversion (50). Engaging in social interactions and adopting approach-oriented behavior might serve as a means to boost baseline dopamine signaling in the cortex. Moreover, the pleasure derived from social encounters could stimulate the reward dopamine system, potentially compensating for lower dopamine levels in synapses.

Furthermore, carriers of the Val allele also exhibit more pronounced self-confidence and ambition. Although Conscientiousness strongly implies volitional control of behavior, leading to the anticipation that carriers of the Met allele might exhibit higher scores, its facets Competence and Achievement Striving are more aligned with approach behavior and the pursuit of rewards rather than with impulse control or delaying reinforcement.

In general, the connections between the *COMT rs4680* polymorphism and personality traits suggest that increased dopamine signaling in carriers of the Met allele is associated with a specific form of sensitivity to negative reinforcement, as well as tendencies toward avoidance behavior and behavioral inhibition. Some previous findings pointed to the increased Neuroticism in Met carriers (e.g. 25,50), which is partially in line with our results. However, our study didn’t show the link to the other aspects of Neuroticism, nor the Neuroticism domain as a whole, suggesting only a specific association with social inhibition. Conversely, lower dopamine levels in Val allele carriers are associated with reward sensitivity, active attitude, and behavioral approach. The expectation of a connection between higher COMT enzyme activity and impulsive behavior, drawn from previous findings (e.g. 17,32), is not supported by the results, indicating that lower dopamine levels are associated with approach-oriented behavior rather than uncontrolled impulsivity.

### DNAm in the promoter region of the *MB-COMT* gene and personality traits

Given the significant role of environmental factors in the emergence of epigenetic modifications, the study also focused on exploring how these changes correlate with personality traits. The objective was to investigate the link between DNAm levels in CpG sites of the *COMT* promoter and personality trait expression, aiming to contribute to the understanding of the mechanisms through which environmental factors affect personality phenotypes. Findings indicate that DNAm of certain CpG sites of the *MB-COMT* promoter has additional effects on dopamine function and consequently behavior, revealing a specific pattern of associations. Notably, traits that are more pronounced in individuals with higher methylated CpG sites of the *MB-COMT* gene promoter are quite distinct from traits that are less pronounced in the case of elevated DNAm. Furthermore, a pattern emerges wherein methylated CpG sites associated with heightened levels of certain personality traits differ from those associated with lower levels of some traits. Hence, it appears that DNAm at different sites on the *MB-COMT* gene promoter may not uniformly affect behavior.

Traits more pronounced in individuals with higher DNAm levels are primarily linked to CpG1, CpG2, and CpG5 sites. These include facets of Extraversion (Gregariousness, Warmth) and Agreeableness (Trust, Modesty), which foster quality relationships and intimacy, suggesting that DNAm at certain sites, mostly CpG2, of the *MB-COMT* gene promoter uniquely impact traits associated with affiliation potential through its influence on dopaminergic function. Moreover, higher DNAm of CpG1, CpG3, and CpG5 is associated with greater sociability, increased openness to values and tolerance for diversity, as well as heightened modesty. Generally, traits more pronounced in individuals with higher methylated CpGs are those relevant to smooth social interactions and harmonious relationships: friendly demeanor, trust in others, acceptance of diversity, and modesty. Since increased DNAm is predominantly associated with gene silencing, it appears that the enhanced expression of these traits results from the lower amounts of COMT enzyme, leading to heightened dopamine signaling in some parts of the brain.

On the other hand, a much larger number of traits, across all five personality domains, seems to be less pronounced in individuals with higher levels of DNAm. These include some of the domains mentioned before, but now associated with DNAm levels at different sites: CpG2, CpG3, and CpG5. For example, some facets of Extraversion were found to be associated with CpG1 and CpG2 sites under the “greater” alternative, while the lower expression of other facets of the same domain are associated with higher levels of DNAm at the CpG5 site including Excitement Seeking, Positive Emotion, and Extraversion itself. Similarly, some of the Agreeableness facets (Trust, Modesty) were associated with CpG2 and CpG5 sites under the “greater” alternative, while others (Straightforwardness, Tender-Mindedness) are associated with CpG3 and CpG4 sites under the “less” alternative.

Several facets of Neuroticism exhibit lower expression in individuals with higher DNAm levels. Most notably, lower Hostility is associated with higher DNAm at CpG2, CpG3, and CpG5. This finding indicates the association between an angry and hostile attitude and decreased dopamine signaling, potentially leading to a sense of reward deprivation. Namely, tendency to frequently experience anger may partly originate from the perception of being unfairly treated (52) and from the belief that others receive more reinforcement than oneself. It may imply somewhat decreased control as well.

Previous research has linked impulsivity to the activity of the *COMT* gene (17,32,41). Our findings show that Impulsiveness is negatively related to the DNAm level of CpG5, if the *MB-COMT* gene is silenced and dopamine degradation function declines, Impulsiveness (N) is lower. This finding aligns with expectations, as Impulsiveness entails a weakened control over basic urges and an immediate quest for reinforcement, which is possibly related to reduced dopamine levels in certain cortical regions. A somewhat similar explanation applies to Excitement Seeking and Positive Emotions facets within the Extraversion domain, which also exhibit decreased expression in individuals with higher DNAm levels at CpG5, and to the association of Openness and CpG4 DNAm as well. Here, lower dopamine signaling seems to drive the quest for reward and positive reinforcement. While Impulsiveness, as the Neuroticism facet, involves low self-control, heightened tension (4) and the pursuit of means to reduce it, Excitement-Seeking and Positive Emotions simply involve continual searching for positive stimulation. Nevertheless, these dimensions all seem to imply some form of compensating for the deficiency of dopamine in the system through the pursuit of reward. Moreover, the association between Straightforwardness (A) and DNAm levels of CpG3 and CpG4 further supports the link between dopamine and behavior control, as honesty and simplicity may imply some form of unrestrained behavior, albeit differing in nature from Impulsiveness. Although not associated with the lack of control, the Agreeableness domain and its Tender-Mindedness facet may be linked to decreased dopamine levels through heightened sensitivity to reward signals, as these traits facilitate positive social reinforcement and acceptance.

Findings regarding the association between depression and COMT gene polymorphism are inconsistent (53), suggesting that the relationship might be complex. Furthermore, epigenetic studies suggest different effects of DNAm in the promoter of *MB-COMT* on prefrontal white matter connectivity in healthy individuals compared to those with major depressive disorder (MDD). While higher *MB-COMT* DNAm is associated with weaker connectivity in healthy subjects, it’s linked with improved connectivity in MDD patients (54). Our results indicate a negative correlation between Depressiveness and increased CpG5 DNAm, suggesting that decreased dopamine levels might lead to a propensity for frequent experiences of sadness and diminished motivation. This association was somewhat surprising, particularly given its alignment with the connection to Positive Emotion. One potential explanation involves a sense of being deprived of rewards due to decreased dopamine signaling, which could trigger feelings of anger, as demonstrated in the case of Angry Hostility, but also feelings of sadness and decline in motivation (55). Additionally, depression is associated with reduced cognitive control, which may result from motivational deficits (56). Furthermore, the findings that individuals with high DNAm levels in CpG5 and CpG4 score lower on the Conscientiousness facets of Order and Competence suggest that traits associated with organization and preparedness could also stem from attempts to activate the reward system in response to reduced dopamine levels, but possibly to overcompensate the sense of insufficient behavioral control as well.

The findings obviously indicate a variety of behavioral responses to decreased dopamine levels in the reward system and suggest the likely role of numerous mechanisms, many of them beyond DNAm of *MB-COMT* gene, that may contribute to shaping these responses. For example, some of these behavioral traits imply a lack of control, and the others possibly an effort to maintain control. It is crucial to investigate environmental factors that contribute to decreasing or increasing DNAm, consequently leading to behavioral changes. Some of these factors, for example, could involve negative life experiences and stress (57), as is possibly the case with Depression and Angry Hostility.

### Are polymorphisms of the *MB-COMT* gene associated with epigenetic changes?

Although the small sample in the epigenetic aspect of the study made it impossible to examine the relationships of specific polymorphisms with DNAm levels, indirectly one can observe tendencies that are important for tracing future research questions. Namely, the findings regarding the relationship between DNAm status at specific CpG sites of the *MB-COMT* promoter and the expression of personality traits suggest a more complex role of this gene in behavior, which is influenced by environmental factors. Notably, the observed correlation between increased DNAm at CpG1 and heightened Gregariousness, and CpG2 DNAm with Warmth, appears somewhat at odds with findings indicating increased expression of these traits in Val allele carriers, given that higher DNAm typically suppresses gene expression. Furthermore, higher CpG2 DNAm was found to have close to significant correlation with higher Competence, whereas CpG4 DNAm is associated with lower Competence. This implies that DNAm at particular CpG sites of the *MB-COMT* gene exerts additional effects on dopamine function and behavior, possibly interacting with *COMT* polymorphism to influence behavior (58). Since epigenetic changes are dynamic and influenced by environmental factors such as effort and stress (59), it’s also possible that some transient factors contributed to these results.

## Limitations and Future Directions

The key strengths of this study lie in using the methodological advantages of twin samples to explore the interplay between genetic factors and environmental influences on personality traits. However, certain limitations warrant consideration, potentially hindering straightforward generalization of the findings. Firstly, the use of a SNP approach in the GWAS era may oversimplify the interpretation of results. Namely, psychological phenotypic features, which are the focus of this study, are determined by a multitude of genes and their possible interactions. Therefore, our findings should be regarded as a contribution to the growing body of evidence linking the *COMT* gene to behaviors associated with impulse control and reward seeking. Secondly, the sample of twins in the epigenetic part of the study is relatively modest. Although the statistical methodologies employed were tailored to address the specific research inquiries, replication on larger cohorts is warranted for robust validation. Thirdly, the hierarchical nature of the psychological phenotypes examined in this study, such as personality traits, underscores the importance of instrument reliability in their assessment. Despite potential limitations in the reliability of individual facets, the NEO-PI-R is widely used measure of the five-factor model of personality, demonstrating validity in prior studies with Serbian twin samples (60,61) and facilitating cross-cultural and cross-linguistic comparisons (12).

## Conclusions

The study reveals significant links between the Met158Val polymorphism of the *COMT* gene and personality traits like Neuroticism, Extraversion, and Conscientiousness. Met allele carriers tend to have higher self-consciousness and social anxiety, while Val allele carriers exhibit more extraversion, sociability, and ambition. This suggests that *COMT* has pleiotropic effects, influencing dopamine signaling and behavior, impacting approach and avoidance tendencies.

Exploring DNAm levels of CpG sites in the *MB-COMT* gene’s promoter region shows a complex relationship with personality traits. Higher DNAm levels at certain CpG sites are linked to traits like warmth, trust, and sociability, while lower DNAm levels relate to reduced expression of traits like hostility and impulsivity. These findings underscore how epigenetic modifications affect dopamine function and behavior, necessitating further study on their interaction with genetic and environmental factors.

Although the study didn’t directly examine the link between specific polymorphisms and DNAm levels in *MB-COMT*, indirect observations suggest a complex interplay between genetic variation and epigenetic modifications in shaping behavior. This emphasizes the importance of considering both genetic and epigenetic factors to understand individual differences in personality and behavior, paving the way for future research on gene-environment interactions.

## Declarations of interest

The authors report no conflict of interest.

## Funding statement

This research was fully supported by the Science Fund of the Republic of Serbia (#7744418, Genetic and environmental influences on psychological adaptation of children and adults – GENIUS).

## Author Contributions

**Dušanka Mitrović:** Conceptualization, Investigation, Methodology, Writing - Original Draft Preparation; **Snežana Smederevac:** Conceptualization, Investigation, Methodology, Writing - Original Draft Preparation, Funding acquisition; **Lissette Delgado-Cruzata:** Investigation, Resources, Supervision, Writing - Review & Editing; **Selka Sadiković:** Conceptualization, Data Curation, Investigation, Methodology, Formal analysis, Writing - Original Draft Preparation; **Dejan Pajić:** Data Curation, Investigation, Methodology, Formal analysis, Software, Visualization, Writing - Original Draft Preparation; **Mechthild Prinz**: Investigation, Resources, Supervision, Writing - Review & Editing; **Zoran Budimlija**: Investigation, Resources, Supervision, Writing - Review & Editing; **Milan Oljača:** Data Curation, Investigation, Methodology, Writing - Review & Editing; **Jelena Kušić Tišma:** Data Curation, Investigation, Writing - Review & Editing; **Nataša Vučinić:** Data Curation, Investigation, Writing - Review & Editing; **Aleksandra Milutinović:** Data Curation, Investigation, Writing - Review & Editing; All authors provided critical feedback, contributed to manuscript editing, and approved the final version of the manuscript.

## Supplementary material

**Table A.** Descriptive statistics by allele groups for the COMT gene on FFM personality domains and facets: molecular genetic sample.

|  | *COMT* | *M* | *SD* |
| --- | --- | --- | --- |
| Neuroticism (trait) | Met+ | 86.67 | 15.90 |
|  | Val+ | 83.48 | 17.94 |
| Anxiety | Met+ | 16.54 | 3.42 |
|  | Val+ | 15.95 | 3.81 |
| Hostility | Met+ | 13.53 | 4.70 |
|  | Val+ | 12.84 | 4.91 |
| Depressiveness | Met+ | 13.23 | 4.06 |
|  | Val+ | 12.66 | 4.07 |
| Self-consciousness | Met+ | 14.99 | 4.17 |
|  | Val+ | 13.98 | 4.17 |
| Impulsivity | Met+ | 16.89 | 4.01 |
|  | Val+ | 17.27 | 4.60 |
| Vulnerability | Met+ | 11.50 | 4.27 |
|  | Val+ | 10.78 | 4.47 |
| Extraversion | Met+ | 114.19 | 19.69 |
|  | Val+ | 119.77 | 16.70 |
| Warmth | Met+ | 21.55 | 4.75 |
|  | Val+ | 23.22 | 4.33 |
| Gregariousness | Met+ | 19.57 | 4.82 |
|  | Val+ | 21.25 | 4.57 |
| Assertiveness | Met+ | 15.65 | 4.24 |
|  | Val+ | 15.82 | 4.07 |
| Activity | Met+ | 18.55 | 4.68 |
|  | Val+ | 19.38 | 4.39 |
| Excitement seeking | Met+ | 18.22 | 4.89 |
|  | Val+ | 18.37 | 5.25 |
| Positive emotions | Met+ | 20.65 | 3.87 |
|  | Val+ | 21.72 | 3.57 |
| Openness to experience (trait) | Met+ | 117.89 | 15.36 |
|  | Val+ | 117.42 | 14.83 |
| Fantasy | Met+ | 19.43 | 4.67 |
|  | Val+ | 19.06 | 4.41 |
| Esthetics | Met+ | 21.44 | 4.19 |
|  | Val+ | 20.75 | 4.45 |
| Openness to feelings | Met+ | 21.60 | 3.58 |
|  | Val+ | 21.66 | 4.05 |
| Openness to action | Met+ | 16.92 | 3.66 |
|  | Val+ | 17.57 | 4.15 |
| Openness to ideas | Met+ | 19.39 | 4.96 |
|  | Val+ | 19.42 | 5.00 |
| Openness to values | Met+ | 19.11 | 3.38 |
|  | Val+ | 18.96 | 3.69 |
| Conscientiousness (trait) | Met+ | 124.88 | 17.25 |
|  | Val+ | 127.38 | 17.61 |
| Competence | Met+ | 21.91 | 3.53 |
|  | Val+ | 22.89 | 3.96 |
| Order | Met+ | 17.78 | 4.06 |
|  | Val+ | 17.40 | 4.10 |
| Dutifulness | Met+ | 24.71 | 3.57 |
|  | Val+ | 24.63 | 3.53 |
| Achievement striving | Met+ | 21.61 | 5.16 |
|  | Val+ | 22.84 | 4.29 |
| Self-discipline | Met+ | 20.72 | 4.44 |
|  | Val+ | 21.19 | 4.41 |
| Deliberation | Met+ | 18.16 | 5.06 |
|  | Val+ | 18.44 | 5.41 |

**Table B.**
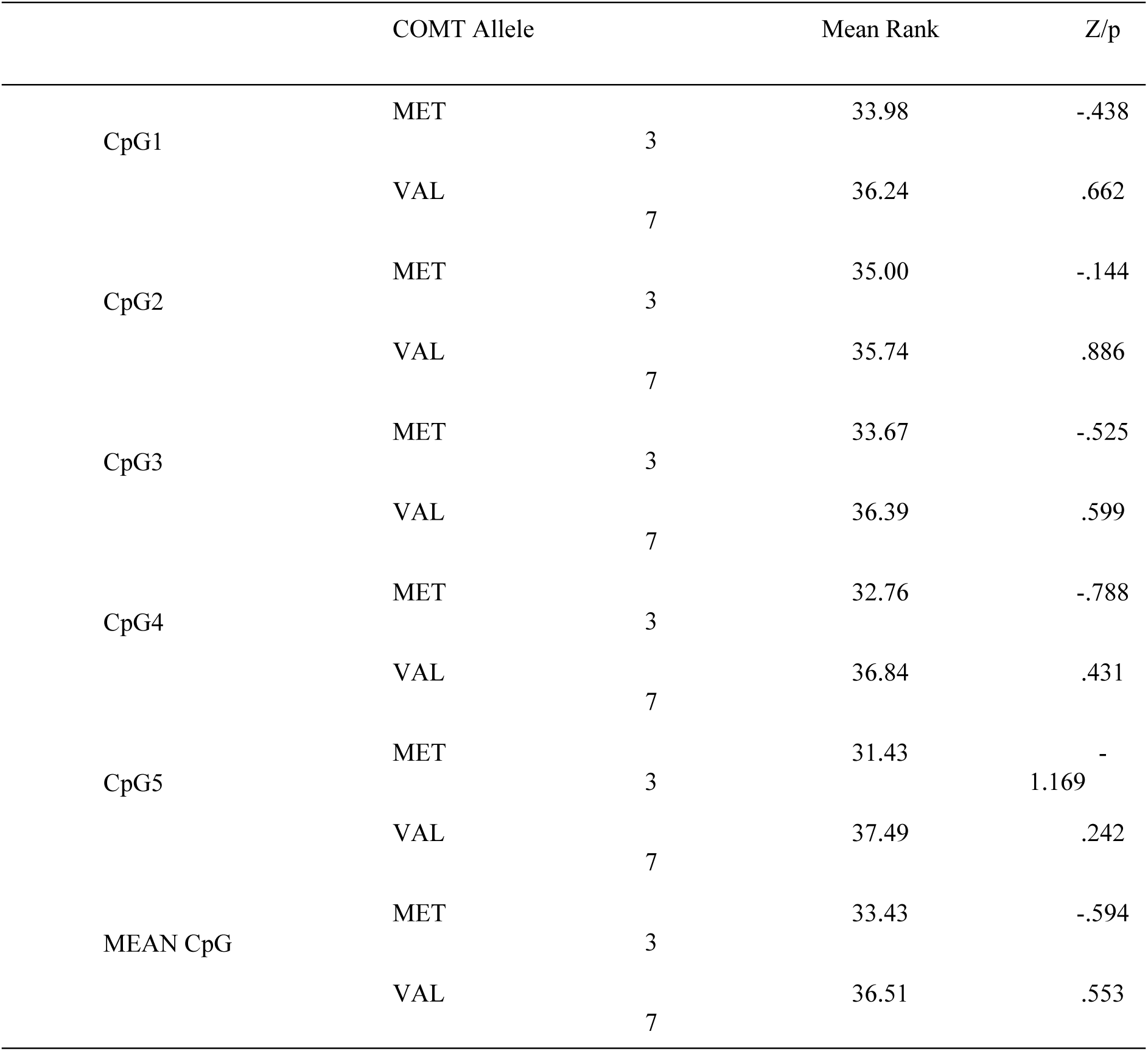
Main effects of COMT alleles on CpG.

**Table C.** Association between COMT DNAm levels and NEO-PI-R dimensions and facets (“greater” alternative)

| Dimension/Facet | CpG1 |  | CpG2 |  | CpG3 |  | CpG4 |  | CpG5 |  |
| --- | --- | --- | --- | --- | --- | --- | --- | --- | --- | --- |
|  | BT | p | BT | p | BT | p | BT | p | BT | p |
| Neuroticism | -0.524 | 1.000 | 0.364 | 0.388 | -0.277 | 1.000 | 0.784 | 0.299 | -1.148 | 1.000 |
| Anxiety | 0.828 | 0.330 | 0.411 | 0.370 | 1.690 | 0.062 | 1.394 | 0.11 | 0.277 | 0.440 |
| Hostility | -0.174 | 1.000 | -1.012 | 1.000 | -0.552 | 1.000 | -0.195 | 1.000 | -1.513 | 1.000 |
| Depression | -0.503 | 1.000 | 1.012 | 0.240 | 0.552 | 0.356 | -0.488 | 1.000 | -1.203 | 1.000 |
| Self-Consciousness | 0.155 | 0.480 | 0.364 | 0.388 | 0.416 | 0.392 | 0.098 | 0.525 | -1.148 | 1.000 |
| Impulsiveness | 0.122 | 0.482 | 0.864 | 0.243 | -2.035 | 1.000 | -0.103 | 1.000 | -0.368 | 1.000 |
| Vulnerability | -1.513 | 1.000 | -0.733 | 1.000 | 0.277 | 0.442 | 0.588 | 0.325 | -0.895 | 1.000 |
| Extraversion | 0.488 | 0.342 | 0.733 | 0.314 | 0.000 | 1.000 | -0.296 | 1.000 | -2.155 | 1.000 |
| Warmth | 0.195 | 0.481 | **2.035** | **0.030** | 0.000 | 1.000 | -1.085 | 1.000 | 0.784 | 0.306 |
| Gregariousness | **1.855** | **0.036** | 1.109 | 0.199 | 0.697 | 0.326 | -0.296 | 1.000 | -0.784 | 1.000 |
| Assertiveness | 0.879 | 0.211 | 1.343 | 0.102 | 0.697 | 0.326 | 0.296 | 0.432 | -0.588 | 1.000 |
| Activity | 1.179 | 0.145 | 1.374 | 0.114 | 1.519 | 0.081 | -0.195 | 1.000 | -0.155 | 1.000 |
| Excitement-Seeking | -1.179 | 1.000 | -1.053 | 1.000 | -0.138 | 1.000 | -0.488 | 1.000 | -2.56 | 1.000 |
| Positive Emotions | -0.138 | 1.000 | 1.331 | 0.106 | -0.986 | 1.000 | -1.394 | 1.000 | -2.355 | 1.000 |
| Openness to Experience | 0.524 | 0.327 | 1.343 | 0.102 | 0.970 | 0.231 | -2.155 | 1.000 | -0.895 | 1.000 |
| Fantasy | -1.179 | 1.000 | 0.324 | 0.419 | 0.552 | 0.356 | -1.172 | 1.000 | -1.203 | 1.000 |
| Aesthetics | -0.828 | 1.000 | -1.497 | 1.000 | -0.282 | 1.000 | -2.092 | 1.000 | -0.277 | 1.000 |
| Feelings | -0.503 | 1.000 | 1.374 | 0.114 | 1.243 | 0.125 | -1.172 | 1.000 | 0.834 | 0.274 |
| Actions | 0.879 | 0.211 | 0.986 | 0.199 | 0.697 | 0.326 | 0.296 | 0.432 | 0.098 | 0.492 |
| Ideas | 0.174 | 0.477 | 0.324 | 0.419 | 0.552 | 0.356 | -1.172 | 1.000 | -1.203 | 1.000 |
| Values | 1.513 | 0.090 | 1.425 | 0.088 | **1.801** | **0.045** | 1.469 | 0.090 | 1.576 | 0.088 |
| Agreeableness | -1.179 | 1.000 | 0.000 | 1.000 | -0.828 | 1.000 | -2.538 | 1.000 | -0.524 | 1.000 |
| Trust | 0.220 | 0.433 | **2.745** | **0.004** | 1.571 | 0.084 | 0.404 | 0.388 | 0.742 | 0.356 |
| Straightforwardness | -0.851 | 1.000 | -0.324 | 1.000 | -1.933 | 1.000 | -2.929 | 1.000 | -0.834 | 1.000 |
| Altruism | 0.834 | 0.299 | 0.733 | 0.314 | 1.108 | 0.158 | -0.588 | 1.000 | -1.148 | 1.000 |
| Compliance | -0.503 | 1.000 | -0.364 | 1.000 | 0.552 | 0.356 | 0.195 | 0.468 | 0.834 | 0.274 |
| Modesty | 0.503 | 0.368 | -0.324 | 1.000 | -1.243 | 1.000 | 0.488 | 0.368 | **1.882** | **0.035** |
| Tender-Mindedness | -1.519 | 1.000 | -0.083 | 1.000 | 0.423 | 0.368 | -1.394 | 1.000 | -1.662 | 1.000 |
| Conscientiousness | 1.172 | 0.210 | 0.733 | 0.314 | -0.697 | 1.000 | -1.676 | 1.000 | -0.784 | 1.000 |
| Competence | 0.524 | 0.327 | 1.701 | 0.054 | -0.416 | 1.000 | -1.469 | 1.000 | -0.214 | 1.000 |
| Order | 0.195 | 0.481 | -0.733 | 1.000 | -0.697 | 1.000 | -1.085 | 1.000 | -1.959 | 1.000 |
| Dutifulness | 0.828 | 0.330 | -0.287 | 1.000 | -1.127 | 1.000 | -2.092 | 1.000 | -1.108 | 1.000 |
| Achievement Striving | 0.851 | 0.274 | 1.012 | 0.240 | 0.552 | 0.356 | -1.172 | 1.000 | -1.203 | 1.000 |
| Self-Discipline | 0.503 | 0.368 | 0.687 | 0.304 | -1.243 | 1.000 | 0.488 | 0.368 | -0.155 | 1.000 |
| Deliberation | 0.250 | 0.463 | 0.222 | 0.435 | 1.041 | 0.169 | -0.210 | 1.000 | 0.710 | 0.263 |

Notes: BT – Barnard’s test, p – p-value for the test. Significant associations are bolded.

**Table D.**
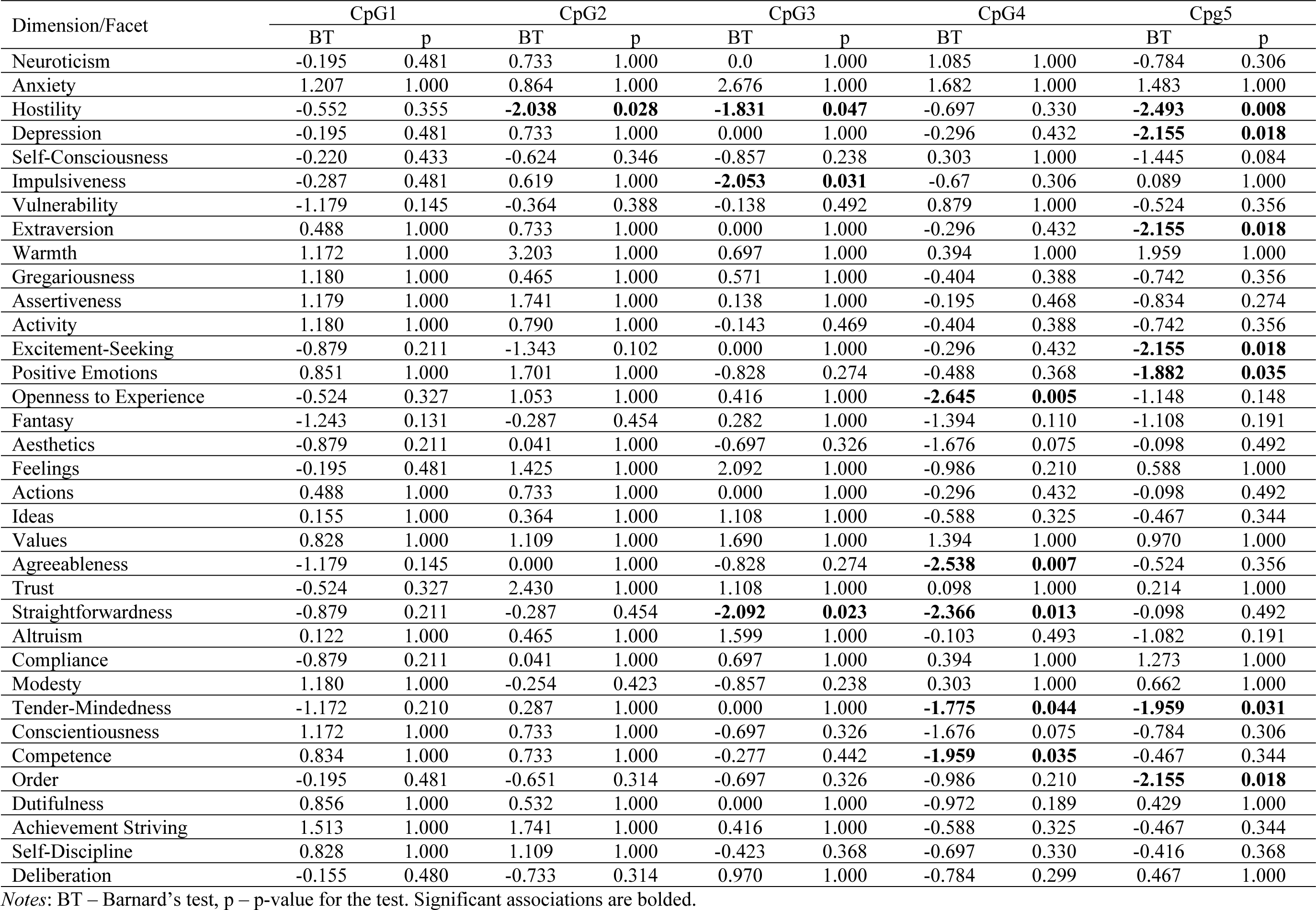
Association between COMT DNAm levels and NEO-PI-R dimensions and facets (“less” alternative)

## Notes

### Competing Interest Statement

The authors have declared no competing interest.

## References

1. Jones PA. Functions of DNA methylation: islands, start sites, gene bodies and beyond. Nat Rev Genet. 2012 Jul;13(7):484–92.

2. Gescher DM, Kahl KG, Hillemacher T, Frieling H, Kuhn J, Frodl T. Epigenetics in Personality Disorders: Today’s Insights. Front Psychiatry. 2018 Nov 19;9.

3. Thomas N, Gurvich C, Kulkarni J. Borderline personality disorder, trauma, and the hypothalamus-pituitary-adrenal axis. NDT. 2019 Sep 9;15:2601–12.

4. Costa PT Jr, McCrae RR. Revised NEO Personality Inventory (NEO-PI-R) and NEO Five-Factor Inventory (NEO-FFI) professional manual. Odessa, FL: Psychological Assessment Resources; 1992.

5. Moore M, Schermer JA, Paunonen SV, Vernon PA. Genetic and environmental influences on verbal and nonverbal measures of the Big Five. Personality and Individual Differences. 2010 Jun 1;48(8):884–8.

6. Soto CJ. How Replicable Are Links Between Personality Traits and Consequential Life Outcomes? The Life Outcomes of Personality Replication Project. Psychol Sci. 2019 May 1;30(5):711–27.

7. McCrae RR, Terracciano A, 78 Members of the Personality Profiles of Cultures Project. Universal Features of Personality Traits From the Observer’s Perspective: Data From 50 Cultures. Journal of Personality and Social Psychology. 2005;88(3):547–61.

8. Jang KL, Livesley WJ, Ando J, Yamagata S, Suzuki A, Angleitner A, et al. Behavioral genetics of the higher-order factors of the Big Five. Personality and Individual Differences. 2006 Jul 1;41(2):261–72.

9. Yamagata S, Suzuki A, Ando J, Ono Y, Kijima N, Yoshimura K, et al. Is the genetic structure of human personality universal? A cross-cultural twin study from North America, Europe, and Asia. Journal of Personality and Social Psychology. 2006;90(6):987–98.

10. Jang KL, Livesley WJ, Angleitner A, Riemann R, Vernon PA. Genetic and environmental influences on the covariance of facets defining the domains of the five-factor model of personality. Personality and Individual Differences. 2002 Jul 5;33(1):83– 101.

11. Sadiković S. Bihevioralnogenetičke osnove petofaktorskog modela ličnosti. [Behavioral genetic basis of the Five-factor Model of Personality] [Internet] [PhD Thesis]. [Novi Sad]: University of Novi Sad, Faculty of Philosophy; 2023. Available from: https://nardus.mpn.gov.rs/bitstream/handle/123456789/21533/Disertacija_13886.pdf?sequence=1

12. Smederevac S, Mitrović D, Sadiković S, Riemann R, Bratko D, Prinz M, et al. Hereditary and environmental factors of the Five-Factor Model traits: A cross-cultural study. Personality and Individual Differences. 2020 Aug 1;162:109995.

13. Terracciano A, Tanaka T, Sutin AR, Deiana B, Balaci L, Sanna S, et al. BDNF Val66Met is Associated with Introversion and Interacts with 5-HTTLPR to Influence Neuroticism. Neuropsychopharmacol. 2010 Apr;35(5):1083–9.

14. Paunonen SV, Ashton MC. Big Five factors and facets and the prediction of behavior. Journal of Personality and Social Psychology. 2001;81(3):524–39.

15. Kandler C, Riemann R, Spinath FM, Angleitner A. Sources of Variance in Personality Facets: A Multiple-Rater Twin Study of Self-Peer, Peer-Peer, and Self-Self (Dis)Agreement. Journal of Personality. 2010;78(5):1565–94.

16. Soto CJ, John OP. The next Big Five Inventory (BFI-2): Developing and assessing a hierarchical model with 15 facets to enhance bandwidth, fidelity, and predictive power. Journal of Personality and Social Psychology. 2017;113(1):117–43.

17. Smederevac S, Sadiković S, Čolović P, Vučinić N, Milutinović A, Riemann R, et al. Quantitative behavioral genetic and molecular genetic foundations of the approach and avoidance strategies. Curr Psychol. 2023 Jun 1;42(17):14268–82.

18. Balestri M, Calati R, Serretti A, De Ronchi D. Genetic modulation of personality traits: a systematic review of the literature. International Clinical Psychopharmacology. 2014 Jan;29(1):1.

19. Bratko D, Butković A, Vukasović Hlupić T. Heritability of Personality. Psihologijske teme. 2017 May 9;26(1):1–24.

20. Munafò MR, Flint J. Dissecting the genetic architecture of human personality. Trends in Cognitive Sciences. 2011 Sep 1;15(9):395–400.

21. Turkheimer E. Weak Genetic Explanation 20 Years Later: Reply to Plomin et al. (2016). Perspect Psychol Sci. 2016 Jan 1;11(1):24–8.

22. Wagner J, Orth U, Bleidorn W, Hopwood CJ, Kandler C. Toward an Integrative Model of Sources of Personality Stability and Change. Curr Dir Psychol Sci. 2020 Oct 1;29(5):438–44.

23. Moore AA, Sawyers C, Adkins DE, Docherty AR. Opportunities for an enhanced integration of neuroscience and genomics. Brain Imaging and Behavior. 2018 Aug 1;12(4):1211–9.

24. Haas BW, Smith AK, Nishitani S. Epigenetic Modification of OXTR is Associated with Openness to Experience. Personality Neuroscience. 2018 Jul;1:e7.

25. Chen C, Chen C, Moyzis R, Dong Q, He Q, Zhu B, et al. Sex Modulates the Associations Between the COMT Gene and Personality Traits. Neuropsychopharmacol. 2011 Jul;36(8):1593–8.

26. Peciña M, Mickey BJ, Love T, Wang H, Langenecker SA, Hodgkinson C, et al. DRD2 polymorphisms modulate reward and emotion processing, dopamine neurotransmission and openness to experience. Cortex. 2013 Mar 1;49(3):877–90.

27. DeYoung CG, Cicchetti D, Rogosch FA, Gray JR, Eastman M, Grigorenko EL. Sources of cognitive exploration: Genetic variation in the prefrontal dopamine system predicts Openness/Intellect. Journal of Research in Personality. 2011 Aug 1;45(4):364–71.

28. Smillie LD, Cooper AJ, Proitsi P, Powell JF, Pickering AD. Variation in DRD2 dopamine gene predicts Extraverted personality. Neuroscience Letters. 2010 Jan 14;468(3):234–7.

29. Tsuchimine S, Yasui-Furukori N, Sasaki K, Kaneda A, Sugawara N, Yoshida S, et al. Association between the dopamine D2 receptor (DRD2) polymorphism and the personality traits of healthy Japanese participants. Progress in Neuro-Psychopharmacology and Biological Psychiatry. 2012 Aug 7;38(2):190–3.

30. Demetrovics Z, Varga G, Szekely A, Vereczkei A, Csorba J, Balazs H, et al. Association between Novelty Seeking of opiate-dependent patients and the catechol-*O*-methyltransferase Val158Met polymorphism. Comprehensive Psychiatry. 2010 Sep 1;51(5):510–5.

31. Hoth KF, Paul RH, Williams LM, Dobson-Stone C, Todd E, Schofield PR, et al. Associations between the COMT Val/Met polymorphism, early life stress, and personality among healthy adults. Neuropsychiatric Disease and Treatment. 2006 Jun 1;2(2):219–25.

32. White MJ, Lawford BR, Morris CP, Young RMcD. Interaction Between DRD2 C957T Polymorphism and An Acute Psychosocial Stressor on Reward-Related Behavioral Impulsivity. Behav Genet. 2009 May 1;39(3):285–95.

33. Aluja A, Balada F, Blanco E, Fibla J, Blanch A. Twenty candidate genes predicting neuroticism and sensation seeking personality traits: A multivariate analysis association approach. Personality and Individual Differences. 2019 Apr 1;140:90–102.

34. Valeeva EV, Kashevarov GS, Kasimova RR, Ahmetov II, Kravtsova OA. Association of the Val158Met Polymorphism of the COMT Gene with Measures of Psychophysiological Status in Athletes. Neurosci Behav Physi. 2020 May 1;50(4):485–92.

35. Reuter M, Kuepper Y, Hennig J. Association between a polymorphism in the promoter region of the TPH2 gene and the personality trait of harm avoidance. International Journal of Neuropsychopharmacology. 2007 Jun 1;10(3):401–4.

36. Reuter M, Schmitz A, Corr PJ, Hennig J. Molecular genetics support Gray’s personality theory: The interaction of COMT and DRD2 polymorphisms predicts the behavioral approach system. International Journal of Neuropsychopharmacology. 2006;9:155–66.

37. Tunbridge EM, Narajos M, Harrison CH, Beresford C, Cipriani A, Harrison PJ. Which Dopamine Polymorphisms Are Functional? Systematic Review and Meta-analysis of *COMT*, *DAT*, *DBH*, *DDC*, *DRD1–5*, MAOA, MAOB, TH, VMAT1, and VMAT2. Biological Psychiatry. 2019 Oct 15;86(8):608–20.

38. Lee LO, Prescott CA. Association of the catechol-O-methyltransferase val158met polymorphism and anxiety-related traits: a meta-analysis. Psychiatric Genetics. 2014 Apr;24(2):52.

39. Dreher JC, Kohn P, Kolachana B, Weinberger DR, Berman KF. Variation in dopamine genes influences responsivity of the human reward system. Proceedings of the National Academy of Sciences. 2009 Jan 13;106(2):617–22.

40. Ren Z, Yang W, Qiu J. Neural and genetic mechanisms of creative potential. Current Opinion in Behavioral Sciences. 2019 Jun 1;27:40–6.

41. Smederevac S, Delgado-Cruzata L, Mitrović D, Dinić BM, Bravo TAT, Delgado M, et al. Differences in MB-COMT DNA methylation in monozygotic twins on phenotypic indicators of impulsivity. Front Genet. 2023 Jan 6;13.

42. Smederevac S, Mitrović D, Sadiković S, Milovanović I, Branovački B, Dinić BM, et al. Serbian Twin Registry. Twin Research and Human Genetics. 2019 Dec;22(6):660–6.

43. Costa PT Jr, McCrae RR. Srpska standardizacija NEO petofaktorskog inventara NEO-FFI: forma S [The Serbian standardization of the NEO Five-Factor inventory NEO-FFI: Form S]. Beograd: Sinapsa edicije; 2019.

44. JASP Team. JASP [Internet]. 2022. Available from: https://jasp-stats.org/

45. Cohen J. Statistical power analysis for the behavioral sciences. New York: Lawrence Erlbaum Associates; 1988.

46. Cheng H, He J. Comparison of tests for association of 2 × 2 tables under multiple testing setting. Communications in Statistics - Simulation and Computation. 2023 Jun 3;52(6):2336–48.

47. Virtanen P, Gommers R, Oliphant TE, Haberland M, Reddy T, Cournapeau D, et al. SciPy 1.0: fundamental algorithms for scientific computing in Python. Nat Methods. 2020 Mar;17(3):261–72.

48. Plotly Technologies Inc. Collaborative data science [Internet]. Plotly Technologies Inc.; 2015. Available from: https://plot.ly

49. Chmielowiec J, Chmielowiec K, Suchanecka A, Trybek G, Mroczek B, Małecka I, et al. Associations Between the Dopamine D4 Receptor and DAT1 Dopamine Transporter Genes Polymorphisms and Personality Traits in Addicted Patients. International Journal of Environmental Research and Public Health. 2018 Oct;15(10):2076.

50. Depue RA, Collins PF. Neurobiology of the structure of personality: Dopamine, facilitation of incentive motivation, and extraversion. Behavioral and Brain Sciences. 1999 Jun;22(3):491–517.

51. Stein MB, Fallin MD, Schork NJ, Gelernter J. COMT Polymorphisms and Anxiety-Related Personality Traits. Neuropsychopharmacol. 2005 Nov;30(11):2092–102.

52. Sanz J, García-Vera MP, Magán I. Anger and hostility from the perspective of the Big Five personality model. Scandinavian Journal of Psychology. 2010;51(3):262–70.

53. Antypa N, Drago A, Serretti A. The role of COMT gene variants in depression: Bridging neuropsychological, behavioral and clinical phenotypes. Neuroscience & Biobehavioral Reviews. 2013 Sep 1;37(8):1597–610.

54. Na KS, Won E, Kang J, Kim A, Choi S, Tae WS, et al. Differential effect of COMT gene methylation on the prefrontal connectivity in subjects with depression versus healthy subjects. Neuropharmacology. 2018 Jul 15;137:59–70.

55. Lyon KA, Juhasz G, Brown LJE, Elliott R. Big Five personality facets explaining variance in anxiety and depressive symptoms in a community sample. Journal of Affective Disorders. 2020 Sep 1;274:515–21.

56. Grahek I, Shenhav A, Musslick S, Krebs RM, Koster EHW. Motivation and cognitive control in depression. Neuroscience & Biobehavioral Reviews. 2019 Jul 1;102:371–81.

57. McGowan PO, Sasaki A, D’Alessio AC, Dymov S, Labonté B, Szyf M, et al. Epigenetic regulation of the glucocorticoid receptor in human brain associates with childhood abuse. Nat Neurosci. 2009 Mar;12(3):342–8.

58. van der Knaap LJ, Schaefer JM, Franken IHA, Verhulst FC, van Oort FVA, Riese H. Catechol-O-methyltransferase gene methylation and substance use in adolescents: the TRAILS study. Genes, Brain and Behavior. 2014;13(7):618–25.

59. Wiegand A, Blickle A, Brückmann C, Weller S, Nieratschker V, Plewnia C. Dynamic DNA Methylation Changes in the COMT Gene Promoter Region in Response to Mental Stress and Its Modulation by Transcranial Direct Current Stimulation. Biomolecules. 2021 Nov;11(11):1726.

60. Mitrović D, Mihić L, Sadiković S, Smederevac S. Common genetic and environmental bases of the mental disorders and personality traits: Special focus on the hierarchical model of psychopathology and NEO-PI-R facets. Journal of Personality. 2023;

61. Nikolašević Ž, Dinić BM, Smederevac S, Sadiković S, Milovanović I, Ignjatović VB, et al. Common genetic basis of the five factor model facets and intelligence: A twin study. Personality and Individual Differences. 2021 Jun 1;175:110682.

